# Luminescence-based screening for extracellular vesicle release modulators reveals a role for PI4KIIIβ in exosome biogenesis upon lysosome inhibition

**DOI:** 10.1101/2023.02.23.529257

**Authors:** Maarten P. Bebelman, Caitrin Crudden, Bart Snieder, Evangelia Thanou, Catharina J.M. Langedijk, Margarida Viola, Steven Eleonora, Urszula Baginska, Olaf Cotugno, Jan Paul M. Bebelman, Monique A.J. van Eijndhoven, Leontien Bosch, Ka Wan Li, Martine J. Smit, Guillaume van Niel, August B. Smit, Frederik J. Verweij, D. Michiel Pegtel

## Abstract

Dysregulated extracellular vesicle (EV) release has been implicated in various pathologies, including cancer, neurodegenerative disease and osteoarthritis. Despite clear therapeutic potential, drug screening for EV release modulators has yielded limited success due to the lack of a sensitive and scalable EV read-out system. Here, we employed CRISPR-Cas9 to engineer HEK293 cells expressing HA-NanoLuciferase-(NL)-tagged endogenous CD63. We found that under basal culture conditions, CD63-containing EVs are released via a mechanism that is independent of the exocytic SNARE protein SNAP23, presumably by direct budding from the plasma membrane. Endo-lysosome inhibition by chemical or genetic perturbation of vATPase strongly increased SNAP23 and nSmase2-dependent exosome secretion from intracellular compartments. Proteomic analysis revealed these exosomes are enriched for early- and late endosomal markers, but also for autophagosomal proteins. This suggests that a proportion of these exosomes originate from amphisomes, although chemical inhibition of canonical autophagy did not affect exosome secretion upon lysosome inhibition. Using a broad-spectrum kinase inhibitor screen, we identified and subsequently validated the lipid kinase PI4KIIIβ as a critical mediator of exosome secretion and amphisome-mediated secretory autophagy, upon lysosome inhibition. We conclude that tagging of endogenous CD63 with NanoLuciferase represents a sensitive, scalable reporter strategy that enables identification of (druggable) modulators of EV biogenesis and release under physiological and pathological conditions.

## Introduction

Extracellular vesicles (EVs) are increasingly recognized for their role in intercellular communication. Most, if not all, cell types secrete EVs either by direct budding from the plasma membrane (PM), in which case they are called ectosomes or microvesicles, or by fusion of multivesicular bodies (MVBs) with the PM, then called exosomes, or a combination of the two (Pegtel and Gould, 2019). Exosomes and ectosomes are not easily distinguished by biochemical or physical analysis but both implicated in mediating the horizontal-transport of various cargo molecules, such as proteins, lipids and RNA, between cells (van Niel et al., 2018). As such, EVs partake in short-distance communication, for example by transporting material across the immune-synapse (Mittelbrunn et al., 2011) or between neurons and oligodendrocytes (Fruhbeis et al., 2013), but they have also been found to mediate larger scale inter-organ communication and induce systemic changes (Thomou et al., 2017). Besides their role in homeostasis, it has become clear that EV-mediated communication is deregulated in various pathologies, such as neurodegenerative disorders and cancer (Kalluri and LeBleu, 2020). Cancer cells have been demonstrated to release increased amounts of EVs with oncogenic cargo and pro-tumorigenic properties (Bebelman et al., 2018). Tumor cell EVs contribute to various cancer processes, such as immune-modulation, drug resistance and metastasis.

Despite the clear clinical potential of modulating EV-mediated communication, there are currently only few pharmacological opportunities for their secretion. Inhibition of neutral sphingomyelinase 2-dependent ILV formation using the small molecule GW4869, or interference with Rab27A-mediated docking of MVBs to the PM using the small molecule NexInhib20 are exceptions (Johnson et al., 2016; Luberto et al., 2002; Subramanian et al., 2020; Yang et al., 2018). However, besides a lack of drug-like molecular properties their therapeutic potential is also limited by toxicity (Figuera-Losada et al., 2015; Liu et al., 2021), although a recent study suggests that selective targeting using a NexInhib20 derivative may overcome its toxicity to some extent (Liu et al., 2021). Furthermore, while previous studies have revealed some key components of the EV biogenesis machinery, our knowledge on the mechanisms of EV secretion in health and disease is highly context-dependent and as a consequence far from complete. This lack in knowledge and the limited availability of EV- modulating drugs are a direct consequence of the technical difficulties associated with studying EV secretion. Currently, the main method of EV quantification is based on concentrating EVs from culture supernatant by ultracentrifugation, followed by direct particle measurements, such as nanoparticle tracking analysis, or western blot against EV marker proteins, such as the tetraspanin CD63. This is a time-consuming and labor-intensive process that requires large volumes of culture medium, and thus precludes the screening of larger drug libraries. We previously developed a live cell imaging approach to quantify exosome secretion from single cells (Bebelman et al., 2020). Although this approach eliminates the need for large culture volumes, it is also time-consuming and in its current form, not suitable for large compound or genetic screening methods.

Several approaches have been developed to screen for modulators of EV secretion. Immunocapture of EVs on anti-CD63-coated beads and subsequent FACS-based quantification allowed for targeted short-hairpin RNA (shRNA) screens that helped identify ESCRT components (Colombo et al., 2013) and Rab GTPases (Ostrowski et al., 2010) involved in EV secretion. More recently, a genome-wide CRISPR screening approach enabled the identification of genes involved in EV-mediated miRNA secretion (Lu et al., 2018). Other groups have generated CD63-GFP expressing cell lines and quantified EV secretion directly, by measuring changes in the level fluorescence of the culture supernatant (Im et al., 2019), or indirectly, by measuring the accumulation of cellular CD63- GFP levels as readout for inhibition of EV secretion (Datta et al., 2018). Despite these notable advances, insight into the mechanisms and druggability of EV biogenesis and secretion by these screening methods has been hampered by technical and practical limitations. Recently, the ultrabright NanoLuciferase enzyme (Hall et al., 2012) has been used to create EV reporters that have the potential for robust, fast and scalable quantification of EV secretion compatible with both genetic and pharmacological screens (Bonsergent et al., 2021; Cashikar and Hanson, 2019; Grisard et al., 2022; Gupta et al., 2020; Hikita et al., 2018).

Here we describe the development of HA-NanoLuc-CD63 (HA-NL-CD63) bioluminescent reporter cells using CRISPR/Cas9-technology that allow high-throughput screening for EV modulators.

We used siRNA, CRISPR and chemical perturbations to unravel the EV biogenesis pathways under basal conditions and upon endo-lysosomal dysfunction due to vATPase inhibition. Finally, using a kinase inhibitor library screen, we identified and subsequently validated the kinase PI4KIIIβ as a critical mediator of exosome secretion in cells with dysfunctional endo-lysosomes.

## Results

### Development of a bioluminescent reporter for EV secretion

We designed a bioluminescent reporter for EV secretion by fusing the small and bright NanoLuc luciferase to CD63, a late endosomal tetraspanin that is frequently used as an EV marker protein. NanoLuc is an ultrabright luciferase and therefore we expected that its incorporation into EVs would enable robust EV quantification in culture supernatant without the need for concentrating EVs (Fig. 1a). We placed the NanoLuc sequence at the N-terminus of CD63 to avoid interfering with its C-terminal internalization sequence (Rous et al., 2002) and added a hemagglutinin (HA)-tag to facilitate antibody-based detection of the HA-NanoLuc-CD63 (HA-NL-CD63) fusion protein. During the course of this project, several other groups have developed similar NanoLuc-based EV reporters. A highly similar construct was evaluated by Cashikar and Hanson (Cashikar and Hanson, 2019) who used a lentiviral system for exogenous expression. In line with their findings, we confirm that HA-NL-CD63 traffics to late endosomal compartments (Supplementary Fig. 1a). Furthermore, immuno-electron microscopy reveals that HA-NL-CD63 is correctly sorted into the intraluminal vesicles (ILVs) of MVBs (Fig. 1b, Supplementary Fig. 1b,c).

**Figure 1.**
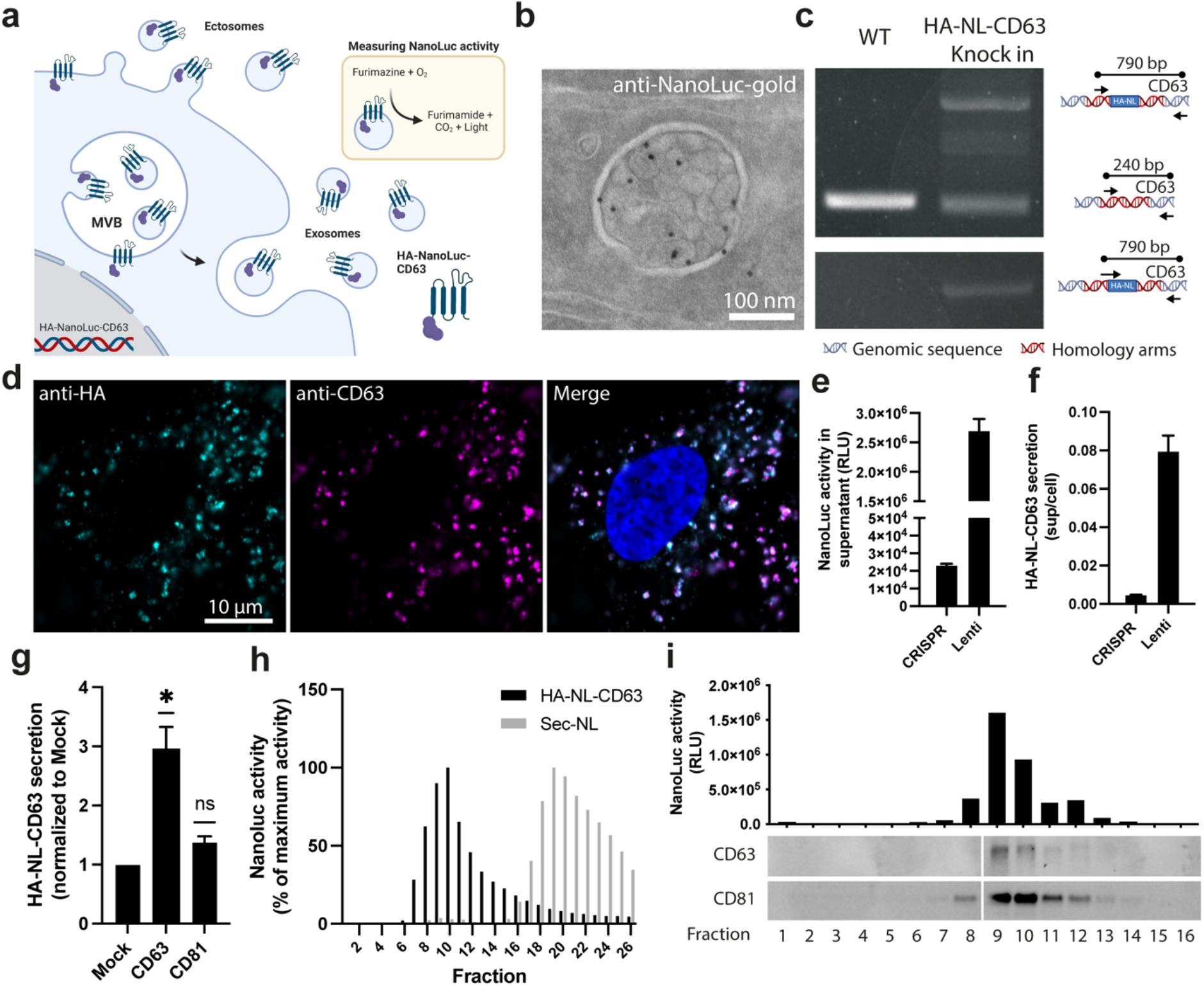
Development and validation of an HA-NanoLuc-CD63 CRISPR knock in cell line. a) Proposed model for the measurement of EV secretion using HA-NL-CD63. This bioluminescent reporter is sorted towards the plasma membrane and multivesicular bodies (MVBs), from where it can be secreted into the extracellular space via ectosomes and exosomes. Direct measurement of bioluminescence in the culture supernatant allows for quantification of CD63-positive EV secretion. Created with BioRender.com. b) EM image of an MVB labelled with gold particles directed to NanoLuc (10 nm) in a HA-NL-CD63– expressing HeLa cell. Magnification from Supplementary Fig. 1b. c) Analysis of HA-NL-CD63 CRISPR knock-in in HEK293 cells by PCR on genomic DNA with primers surrounding the HA-NL insertion site (upper) and with a forward primer within the HA- NL sequence (lower), confirming mono-allelic knock-in of HA-NL. d) Immunofluorescent staining against HA (cyan) and CD63 (magenta) in CRISPR-HA-NL-CD63 HEK293 cells. Blue: DAPI. e,f) Absolute (e) or normalized (f) NanoLuc activity in culture supernatant from equal amounts of CRISPR-HA-NL-CD63 and Lenti-HA-NL-CD63 HEK293 cells. Representative graph from two independent experiments. Data is presented as mean ± S.D. g) HA-NL-CD63 secretion from CRISPR-HA-NL-CD63 HEK293 cells upon transfection with CD63 or CD81. pHluorin-tagged tetraspanin constructs were used to confirm equal transfection efficiency. Data represents mean ± S.E.M. of three independent experiments. ns: non-significant, *P<0.05 using a one- sample, two-tailed T-test. h) NanoLuc activity in supernatant from HEK293 cells endogenously expressing HA-NL-CD63 or transfected with Secreted-NL (Sec-NL) after fractionation by size-exclusion chromatography. Displayed as percentage of maximum activity per reporter. i) NanoLuc activity measurements and western blot on supernatant from HEK293 cells endogenously expressing HA-NL-CD63 after ultrafiltration and Optiprep-density ultracentrifugation.

We then set out to generate stable HA-NL-CD63 expressing cell lines to screen for potential EV production and/or release modulators. We chose HEK293 cells as a cellular background and stably expressed HA-NL-CD63 using lentiviral transduction. Because CD63 itself can drive ILV formation (Edgar et al., 2014; van Niel et al., 2011), exogenous (over)expression of CD63 may affect basal EV biogenesis and secretion. To avoid this, we also tagged endogenous CD63 in HEK293 cells with HA-NanoLuc using CRISPR/Cas9-technology (Fig. 1c,d) and measured NanoLuc activity in the culture supernatant upon addition of the membrane-permeable NanoLuc substrate Furimazine. Both the Lenti-HA-NL-CD63 and the CRISPR-HA-NL-CD63 cell lines secrete HA-NL-CD63 into the culture supernatant. As expected, the NanoLuc activity in the cells and culture supernatant of the CRISPR knock-in cell line was lower than of the lentiviral transduced cell line (Fig. 1e), but sufficient to measure EV secretion in a time-dependent manner (Supplementary Fig. 1d). At sub-confluent cell densities, the cellular NanoLuc activity can be used to normalize the HA-NL-CD63 secretion, correcting for small variations in cell plating (Supplementary Fig. 1e). Importantly, normalized HA-NL-CD63 secretion (NanoLuc activity in supernatant divided by cellular NanoLuc activity) is more than 15-fold higher in Lenti-HA-NL-CD63 cells compared to the CRISPR-HA-NL-CD63 line, suggesting that CD63 overexpression itself strongly stimulates its secretion through EVs (Fig. 1f). Indeed, transient overexpression of CD63, but not CD81, resulted in an increase in fractional release of HA-NL-CD63 (Fig. 1g). In order to avoid exogenous disruption of EV biogenesis, we continued with the CRISPR-HA-NL-CD63 cells for further experiments.

To confirm that the NanoLuc activity in the supernatant is in fact associated with EVs, we fractionated the culture supernatant by size-exclusion chromatography (SEC) and Optiprep density-gradient ultracentrifugation (ODG-UC) and analyzed NanoLuc activity in separate fractions. After SEC, the majority of the NanoLuc activity was detected in the EV-enriched fractions 9 and 10 and near absent from the protein-enriched fractions 18-26, in stark contrast to Sec-NL (NanoLuc enzyme fused to the IL6-secretion peptide) (Fig. 1h). Similarly, after ODG-UC, the NanoLuc activity was present in fractions 9 and 10, which contain the EV-markers CD63 and CD81 (Fig. 1i). Overall, these results show that the CRISPR-HA-NL-CD63 cells faithfully report the secretion of CD63-positive EVs.

### Bafilomycin enhances HA-NL-CD63 secretion by promoting MVB-PM fusion

Previous research by multiple groups, including our own, established involvement of the SNARE molecule SNAP23 in the final secretion step of exosome release from MVBs (Verweij et al., 2018; Wei et al., 2017). To demonstrate that the CRISPR-HA-NL-CD63 cell line can be used to identify modulators of MVB-derived EV secretion, we measured the effect of siRNA-mediated SNAP23 knockdown on HA-NL-CD63 secretion. Although knockdown efficiency was approximately 70%, we did not observe a reduction in HA-NL-CD63 secretion (Supplementary Fig. 2a,b). To ensure complete ablation of the SNAP23 protein, we used CRISPR/Cas9-editing to knock out the SNAP23 gene in the HA-NL-CD63 reporter cells (Supplementary Fig. 2c). Surprisingly, even complete ablation of SNAP23 did not reduce HA-NL-CD63 secretion (Fig. 2a). In fact, both SNAP23 KD and KO increase HA-NL-CD63 secretion. These results suggest that SNAP23 does not contribute to EV-CD63 secretion from HEK293 under basal culture conditions. This agrees with recent reports by others suggesting that under standard culture conditions the majority of EV secretion from HEK293 cells is the result of direct budding from the PM rather than through the fusion of MVBs with the PM (Fordjour et al., 2022).

**Figure 2.**
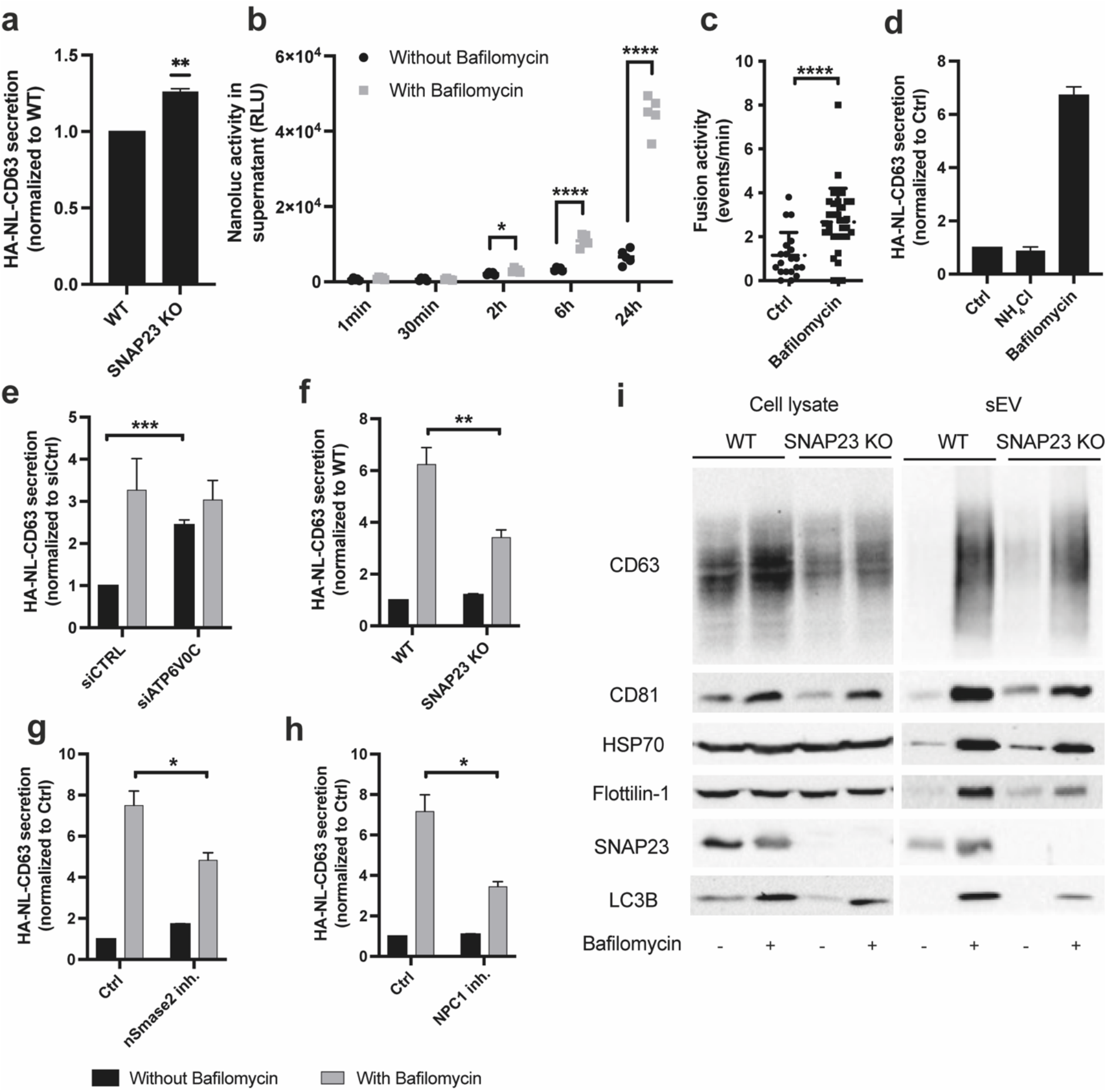
Bafilomycin enhances HA-NL-CD63 secretion by promoting MVB-PM fusion a) Effect of SNAP23 KO on HA-NL-CD63 secretion. Data represents the mean ± S.E.M of three independent experiments. **P<0.01 using a one-sample, two-tailed T test. b) NanoLuc activity in the supernatant of non-treated or bafilomycin-treated CRISPR-HA-NL-CD63 cells at various time points. *p <0.05, ****p <0.0001 using a two-sample, two-tailed Student’s T test. c) Quantification of the fusion of CD63- positive MVBs with the PM in CRISPR-HA-NL-CD63 cells transfected with CD63-pHluorin under basal conditions or upon 3h treatment with 100 nM bafilomycin. d) Effect of NH_4_Cl (20 mM) and bafilomycin (100 nM) on HA-NL-CD63 secretion. Data represents the mean ± S.E.M of three independent experiments. e) Effect of ATP6V0C knockdown on HA-NL-CD63 secretion under basal and bafilomycin-induced conditions. Data represents the mean ± S.E.M of five independent experiments. ***p <0.001 using a one-sample, two-tailed Student’s T test. f) Effect of SNAP23 KO on HA-NL-CD63 secretion under basal and bafilomycin-induced conditions. Data represents the mean ± S.E.M of six independent experiments. **p <0.01 using a two- sample, two-tailed Student’s T test. g) Effect of nSmase2 inhibition (10 μM GW4869) on HA-NL-CD63 secretion under basal and bafilomycin-induced conditions. Data represents the mean ± S.E.M of three independent experiments. *p <0.05 using a two-sample, two-tailed Student’s T test. h) Effect of NPC-1 inhibition (2 μg/ml U18666A) on HA-NL-CD63 secretion under basal and bafilomycin-induced conditions. Data represents the mean ± S.E.M of three independent experiments. *p <0.05 using a two-sample, two-tailed Student’s T test. i) Western blot against EV marker proteins on cell lysates and isolated small EVs from wildtype (WT) and SNAP23 KO HEK293 cells with or without bafilomycin (100 nM) stimulation. Representative of three independent experiments.

In order to identify modulators of exosomes from MVBs specifically, we sought for a strategy to boost MVB-derived exosome secretion from HEK293 cells under identical standard growth conditions. The vacuolar H+ ATPase (vATPase) inhibitor bafilomycin increases exosome secretion in cell types from different origin (Cashikar and Hanson, 2019; Edgar et al., 2016; Guo et al., 2017). In agreement with these reports, we observed a time-dependent increase in HA-NL-CD63 secretion under nanomolar concentrations of bafilomycin treatment (100nM), ranging from a 30% increase compared to non-treated cells at 2h to an 800% increase after 24h of bafilomycin stimulation (Fig. 2b). Probing dose-dependency, we found that a lower concentration of bafilomycin (10 nM) stimulated HA-NL-CD63 secretion to a similar extent (Supplementary Fig. 2d). Consistent with enhanced exosome release from MVBs rather than PM budding, live TIRF imaging with the pH-sensitive CD63-pHluorin reporter (Bebelman et al., 2020) revealed that incubation with bafilomycin for 3h stimulates MVB-PM fusion activity in HEK293 cells (Fig. 2c).

Although the stimulatory effect of bafilomycin on exosome secretion has already been demonstrated by various groups, the underlying mechanism(s) remains unclear. Bafilomycin inhibits the vATPase V0c subunit, thereby increasing the luminal pH of endolysosomes. However, conflicting observations on the role of pH neutralization in bafilomycin-induced exosome secretion have been reported (Cashikar and Hanson, 2019; Guo et al., 2017). To assess this in our CRISPR-based HA-NL- CD63 HEK293 cells, we neutralized acidic compartments with NH_4_Cl and chloroquine and measured the effects on HA-NL-CD63 secretion. NH_4_Cl did not increase HA-NL-CD63 secretion (Fig. 2d) and the increase in secretion upon chloroquine treatment was significantly less (50%), compared to >800% increase upon bafilomycin exposure (Supplementary Fig. 2e). These findings are in full agreement with findings of Cashikar et al. and strongly suggest that in HEK293 cells stimulation of EV secretion by bafilomycin is independent from its effect on endosomal pH.

In addition to vATPase, bafilomycin can inhibit several other ATPases with lower affinity. Bafilomycin has been demonstrated to block autophagosome-lysosome fusion by raising cytosolic calcium levels through inhibition of the sarco/endoplasmic reticulum Ca^2+^-ATPase (SERCA) (Mauvezin et al., 2015). To test whether bafilomycin-induced elevation of intracellular calcium levels may drive EV secretion, we stimulated intracellular calcium levels using the Ca^2+^-ionophore ionomycin and the SERCA inhibitor thapsigargin. Both these compounds did not significantly increase HA-NL-CD63 secretion (Supplementary Fig. 2f,g). Furthermore, the calcium chelator BAPTA-AM did not reduce bafilomycin-induced HA-NL-CD63 secretion (Supplementary Fig. 2h). To confirm that bafilomycin stimulates HA-NL-CD63 secretion through inhibition of the vATPase, and not other ATPases, we next depleted the V0c-subunit of vATPase using siRNA-mediated knockdown. Similar to bafilomycin treatment, knockdown of V0c promotes HA-NL-CD63 secretion, suggesting that bafilomycin stimulates exosome secretion by inhibiting vATPase (Fig. 2e). Using bafilomycin to induce exosome secretion, we assessed the potential of the CRISPR-HA-NL-CD63 cell line to identify modulators of exosome secretion. Notably, knockout of SNAP23 now inhibited ±50% of the bafilomycin-induced HA-NL-CD63 secretion (Fig. 2f). Similarly, inhibition of nSmase2-dependent ILV formation using GW4969 (Trajkovic et al., 2008) decreased bafilomycin-induced, but not basal, HA-NL-CD63 secretion indicating a role for ceramide synthesis (Fig. 2g). Furthermore, inhibition of NPC1, resulting in impaired centrifugal endosomal trafficking and MVB-PM fusion activity in HeLa cells (Verweij et al., 2022), selectively inhibited bafilomycin-induced HA-NL-CD63 secretion (Fig. 2h).

To validate the effect of bafilomycin and SNAP23 depletion on EV secretion with an orthogonal approach, we isolated EVs from bafilomycin-stimulated parental and SNAP23 KO HEK293 cells using a combination of ultrafiltration (UF) and SEC, and performed western blot analysis for various EV markers (Fig. 2i). In line with the HA-NL-CD63 measurements, bafilomycin increased the secretion of the EV markers CD81, Flotillin, Hsp70 in the parental cells, but to a lesser extent in the SNAP23 KO cells. Given recent findings of EV-mediated secretion from amphisomes (Jeppesen et al., 2019), and the fact that bafilomycin treatment blocks the fusion of autophagosomes with lysosomes (Yamamoto et al., 1998), we also assessed the presence of the autophagy marker LC3B in our exosome preparation. Interestingly, bafilomycin stimulates the secretion of LC3B via EVs, and this effect is partially dependent on the expression of SNAP23.

In summary, our results suggest that under basal *in vitro* culture conditions HEK293 cells mainly secrete CD63 via SNAP23/nSmase2-independent mechanisms, presumably by vesicle budding from the PM. Bafilomycin stimulates the secretion of exosomes from MVBs or related organelles through inhibition of vATPase, but independent of endosomal pH neutralization. Finally, the effects of SNAP23 depletion and nSmase2/NPC1 inhibition on bafilomycin-induced HA-NL-CD63 secretion demonstrate that the CRISPR-HA-NL-CD63 cell line can be used to identify proteins involved in exosome biogenesis and release pathways.

### Bafilomycin-induced exosome secretion does not depend on canonical autophagy

Because bafilomycin exposure has been linked to secretory autophagy (Ejlerskov et al., 2013; Solvik et al., 2022), we assessed the effect of bafilomycin on EV cargo proteins by performing mass spectrometry on EVs collected under basal and bafilomycin-induced conditions. Bafilomycin treatment resulted in significant changes in the EV proteome, and we could observe a clear increase in various endosomal proteins, such as CD63, Lamp2, Rab7A, and Rab5 (Fig. 3a,b). GO enrichment analysis using proteins that are present in EVs only under basal conditions, or downregulated upon bafilomycin treatment, showed enrichment of various cellular components associated with the cell surface, such as *cell-substrate junction, focal adhesion, membrane ruffle*, and *cell cortex* (Supplementary Fig. 3a). The same analysis for proteins that are upregulated or present only upon bafilomycin treatment showed enrichment of cellular components associated with the endo- lysosomal system, including *cytoplasmic membrane vesicle, early endosome, late endosome* and *lysosome* (Supplementary Fig. 3b). These findings are consistent with the hypothesis that under basal culture conditions most EVs originate from the PM and that bafilomycin stimulates the secretion of endosome-derived exosomes. Interestingly, gene ontology enrichment analysis revealed enrichment of proteins involved in autophagy (Supplementary Fig. 3c), such as the autophagy receptor SQSTM1/P62 and the ATG8 family members GABARAPL1/2, in the EV pellet from bafilomycin-treated cells (Fig. 3a,b). These results confirm a recent report describing the EV-associated secretion of autophagy receptors upon lysosomal inhibition (SALI) (Solvik et al., 2022). Under SALI conditions, autophagosomes fuse with MVBs, generating hybrid compartments called amphisomes that become competent for fusion with the PM. Notably, the effect of SNAP23 KO on bafilomycin-induced EV-associated LC3B secretion (Fig. 2i) suggests that SNAP23 mediates both MVB-PM and amphisome-PM fusion. Consistent with such a role, SNAP23 KO reduced the total amount of protein in the EV pellet upon bafilomycin treatment, resulting in a lower number of different proteins that could be reliably detected (Fig 3c). However, differential abundance analysis on the proteins that were present in both samples revealed no significant differences between the EV-associated proteomes of bafilomycin- stimulated wildtype (WT) and SNAP23 KO cells (Fig. 3d). This demonstrates that SNAP23 KO affects the secretion of endosome- and autophagy-related proteins equally, indicating that SNAP23 contributes to both exosome secretion and secretory autophagy upon lysosome inhibition.

**Figure 3.**
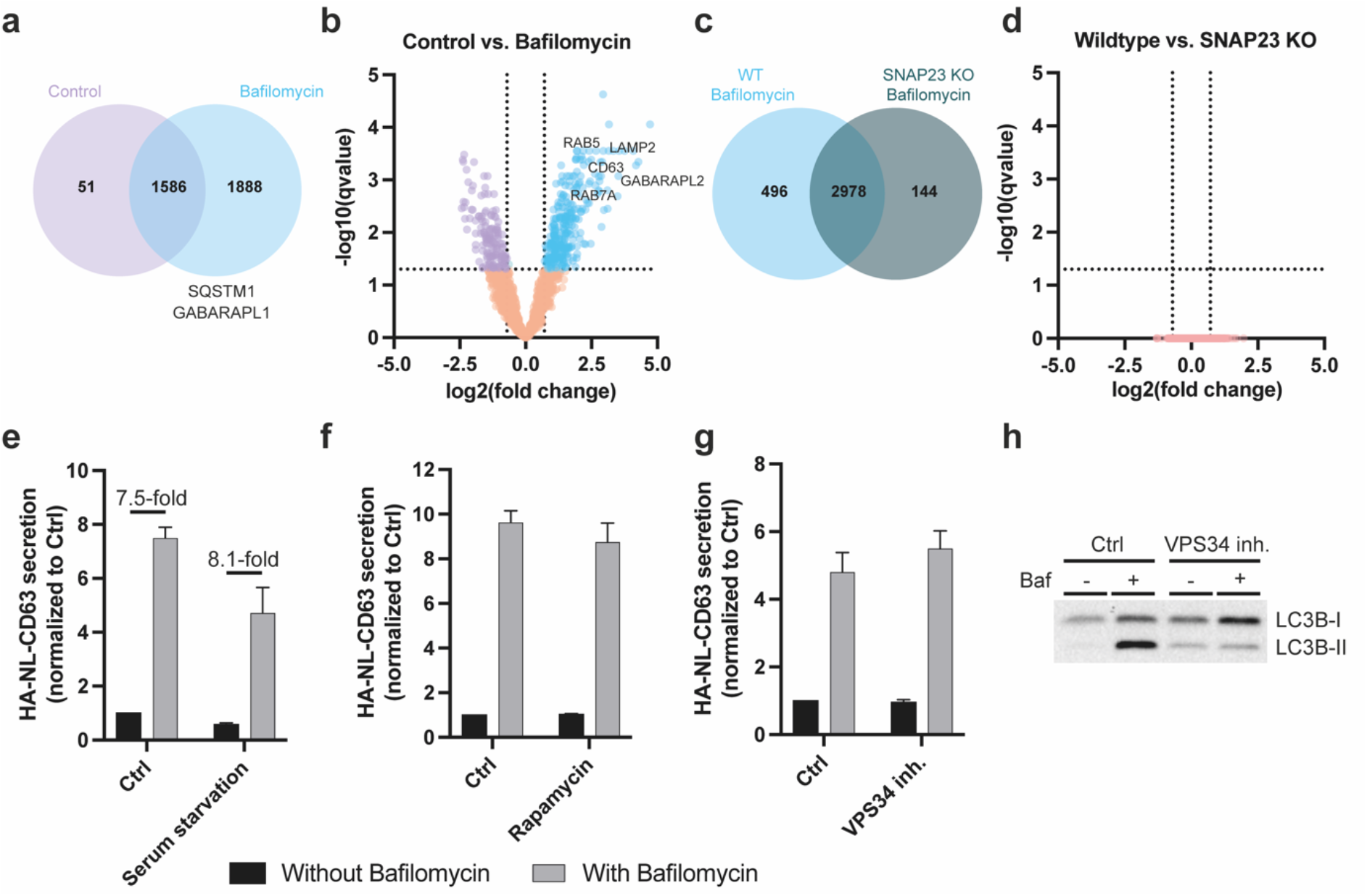
Bafilomycin-induced exosome secretion does not depend on canonical autophagy. a,b) Venn diagram (a) and volcano plot (b) of the proteins in EVs from control and bafilomycin-induced HEK293 cells isolated by UF-SEC-UF. Proteins with a q-value ≤ 0.05, and a log2(fold change) ≤ -0.71 or ≥ 0.71 were considered differentially abundant. Key endo-lysosome- and autophagy-related proteins have been highlighted. c,d) Venn diagram (c) and volcano plot (d) of the proteins in EVs from bafilomycin-induced wildtype (WT) and SNAP23 KO HEK293 cells isolated by UF-SEC-UF. Proteins with a q-value ≤ 0.05, and a log2(fold change) ≤ -0.71 or ≥ 0.71 were considered differentially abundant. e) Effect of serum starvation on HA-NL-CD63 secretion under basal and bafilomycin-induced conditions. Data represents the mean ± S.E.M of three independent experiments. f) Effect of Rapamycin (1 μM) on HA-NL-CD63 secretion under basal and bafilomycin-induced conditions. Data represents the mean ± S.E.M of four independent experiments. g) Effect of VPS34 inhibition (1 μM SAR405) on HA-NL-CD63 secretion under basal and bafilomycin-induced conditions. Data represents the mean ± S.E.M of three independent experiments. h) Western blot for LC3B-I/II demonstrating that 1 μM SAR405 inhibits bafilomycin-induced lipidation of LC3B.

Previous reports have linked autophagy and amphisome formation to changes in exosome secretion (Fader et al., 2008; Hessvik et al., 2016; Peng et al., 2021; Wang et al., 2019). Thus, we questioned whether autophagy and MVB-autophagosome fusion are required to enhance exosome secretion under bafilomycin-stimulated conditions. To investigate this, we stimulated or inhibited canonical autophagy and assessed the effect on HA-NL-CD63 secretion. Stimulation of autophagy by serum starvation reduced both basal and bafilomycin-induced secretion to a similar extent, and thus did not change the relative increase in HA-NL-CD63 secretion upon bafilomycin-treatment (Fig. 3e). Furthermore, induction of autophagy through mTOR-inhibition by rapamycin did not have any effect on HA-NL-CD63 secretion (Fig. 3f). Finally, inhibition of VPS34, a class III phosphoinositide 3-kinase required for autophagosome formation, did not affect basal nor bafilomycin-induced HA-NL-CD63 secretion, despite clearly reduced LC3-lipidation (Fig. 3g,h). We conclude that canonical autophagy is not required for the enhanced secretion of exosomes upon bafilomycin treatment, even though our results corroborate previous findings that lysosomal impairment stimulates exosome-associated secretory autophagy.

### PI4KIIIβ contributes to exosome biogenesis upon lysosome inhibition

To test the potential of our HA-NL-CD63 cells to identify EV production and release modulators in a screening set-up, we performed a broad-spectrum kinase inhibitor screen of 400 small molecule inhibitors for their effect on HA-NL-CD63 secretion in the presence or absence of a non-toxic concentration of bafilomycin (Supplementary Fig. 4a,b). We chose to target kinases as they are frequently used drug targets and multiple kinases have previously been implicated in EV secretion (Imjeti et al., 2017; Wei et al., 2017). We identified 13 kinase inhibitors that significantly decreased basal HA-NL-CD63 secretion and 120 kinase inhibitors that reduced bafilomycin-induced HA-NL-CD63 secretion more than the cut-off value of 2xSD of the DMSO controls. Kinase inhibitors that reduced cellular NanoLuc activity by more than 20% were excluded from further analysis, as we previously observed that a reduction in cellular NanoLuc activity was indicative of decreased cell viability. We compared the effect of kinase inhibitors under bafilomycin-stimulated conditions with their effect on HA-NL-CD63 secretion under basal culture conditions (Fig. 4a). Notably, there was hardly any correlation between the effects of individual kinase inhibitors on HA-NL-CD63 secretion under these two conditions (R^2^ = 0.1773). Amongst the kinase inhibitors that selectively inhibited basal HA-NL-

**Figure 4.**
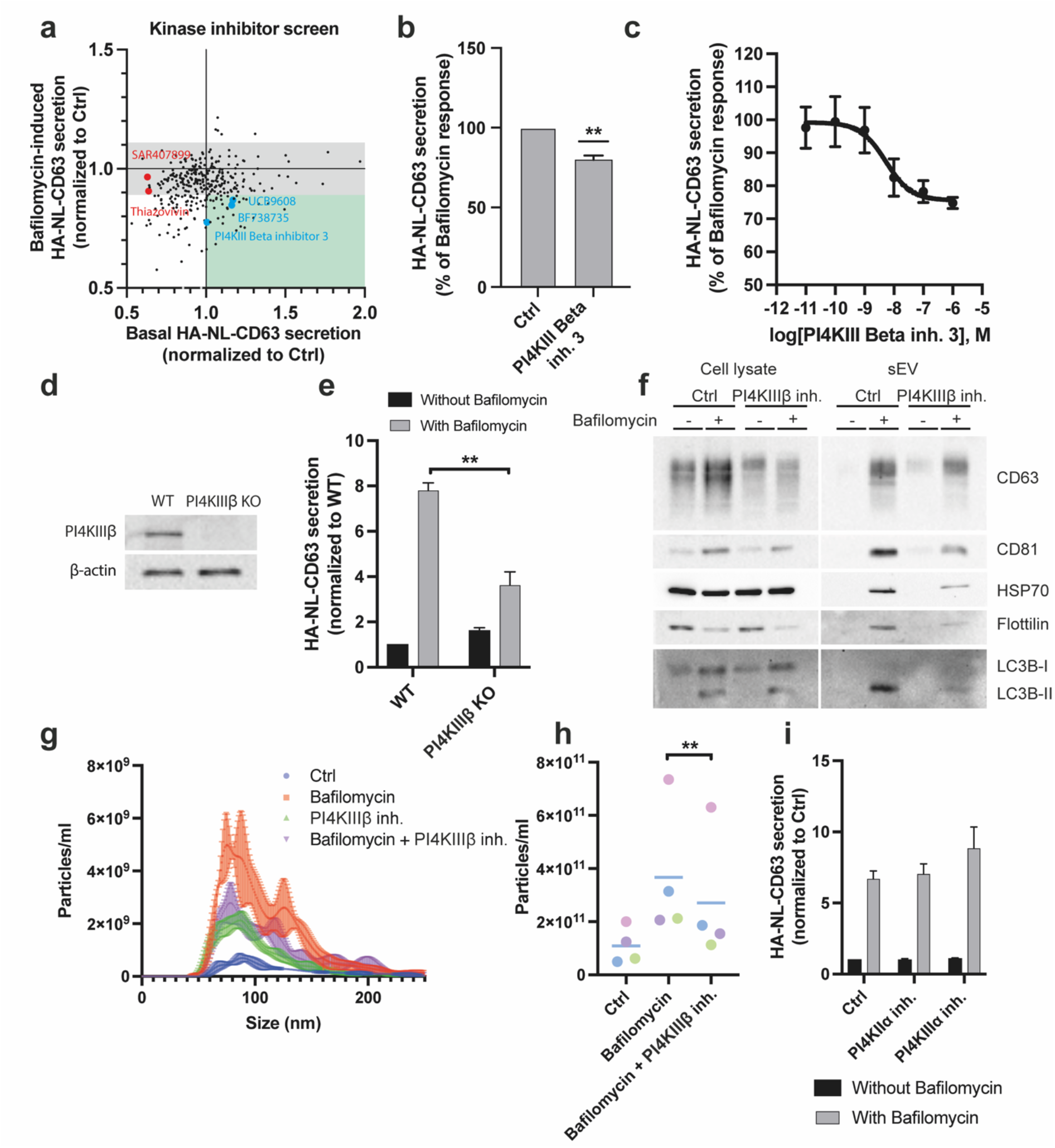
Kinase inhibitor screening reveals PI4KIIIβ as modulator of bafilomycin-induced EV secretion. a) Correlation plot depicting normalized HA-NL-CD63 secretion for individual kinase inhibitors from the kinase inhibitor screen performed under basal and bafilomycin-induced conditions. Grey area corresponds to the mean ± 2 times the standard deviation of the DMSO controls under bafilomycin-induced conditions. Kinase inhibitors in the green area were considered selective inhibitors of bafilomycin-induced HA-NL-CD63 secretion. b) Effect of PI4KIIIβ inhibition (PI4KIII Beta inhibitor 3, 1 μM) on bafilomycin- induced HA-NL-CD63 secretion. Data represents the mean ± S.E.M of four independent experiments. **P<0.01 using a one- sample, two-tailed T-test. c) Dose-inhibition curve of the PI4KIII Beta inhibitor 3 on bafilomycin-induced HA-NL-CD63 secretion. Data is a representative of three independent experiments. The IC_50_ value of 6.4 ±1.5 nM (mean + S.E.M) was determined based on three independent experiments. d) Confirmation of PI4KIIIβ knockout at the protein level in HA-NL- CD63-CRISPR HEK293 cells. e) Effect of PI4KIIIβ KO on HA-NL-CD63 secretion under basal and bafilomycin-induced conditions. Data represents the mean ± S.E.M of three independent experiments. **p <0.01 using a two-sample, two-tailed Student’s T test. f) Western blot against EV marker proteins on cell lysates and isolated small EVs from control and bafilomycin-induced HEK293 cells in the presence or absence of PI4KIIIβ inhibitor (PI4KIII Beta inhibitor 3, 1 μM). Representative of two independent experiments. g) Effect of PI4KIIIβ inhibition (PI4KIII Beta inhibitor 3, 1 μM) on sEV secretion as determined by Nanoparticle Tracking Analysis. Representative of four independent experiments. h) Quantification of (g). Combined data of four independent experiments. Data is color-coded by biological replicate, where dots represent each biological replicate, and the horizontal bars represent mean ± SEM. **, P < 0.01 using a Student’s two-tailed two-sample t-test paired by biological replicate. i) Effect of PI4KIIα (PI-273, 2 μM) and PI4KIIIα (10 nM, GSK-A1) inhibition on HA-NL-CD63 secretion under basal and bafilomycin-induced conditions. Data represents the mean ± S.E.M of three (PI4KIIα) or four (PI4KIIα) independent experiments.

CD63 secretion levels were two inhibitors of ROCK (Fig. 4a, highlighted in red), a kinase that may support vesicle budding from the PM through its role in actomyosin remodelling (Li et al., 2012; Sedgwick et al., 2015). Validation outside of the screen confirmed that the ROCK inhibitors SAR407899 and Thiazovivin inhibited basal, but not bafilomycin-induced EV secretion, which again corroborates the hypothesis that under basal conditions the majority of CD63 secretion occurs at the PM (Supplementary Fig. 4c,d).

Amongst the inhibitors that reduced HA-NL-CD63 secretion specifically under bafilomycin- stimulated conditions were three inhibitors of the lipid kinase phosphatidylinositol 4-kinase III beta (PI4KIIIβ) (Fig. 4a, highlighted in blue). We repeated the HA-NL-CD63 assay for PI4KIII Beta inhibitor 3 outside of the screening set-up and confirmed that it consistently inhibited bafilomycin-stimulated HA-NL-CD63 secretion by around 30% (Fig. 4b). Interestingly, PI4KIIIβ inhibition slightly increased the HA-NL-CD63 secretion under basal (non-bafilomycin treated) conditions (Supplementary Fig. 4e), similar to what we observed for SNAP23 KO and nSmase2 inhibition (Fig. 2f,g). A dose-inhibition curve revealed that PI4KIII Beta inhibitor 3 inhibited bafilomycin-induced secretion with an IC_50_ value of 6.4 ±1.5 nM (mean + S.E.M) (Fig. 4c), which is remarkably close to its reported IC_50_ value of 5.7 nM for PI4KIIIβ (Herman et al., 2013). This strongly suggests that the effect of this inhibitor on HA-NL-CD63 secretion is indeed due to PI4KIIIβ inhibition and not because of non-specific inhibition of other kinases or other off-target effects.

To further validate the role of PI4KIIIβ in bafilomycin-induced HA-NL-CD63 secretion, we used CRISPR-mediated KO of PI4KIIIβ and found that this reduced bafilomycin-induced HA-NL-CD63 secretion by approximately 50% (Fig. 4d,e). We confirmed that PI4KIIIβ inhibits bafilomycin-induced EV secretion using UF-SEC-UF EV isolation followed by western blot analysis for CD63 and several other EV markers, as well as LC3B-II (Fig. 4f). Furthermore, nanoparticle tracking analysis demonstrated that PI4KIIIβ inhibition reduced the bafilomycin-induced secretion of particles in the size-range of small EVs presumably exosomes (Fig. 4g,h). To show these findings are not HEK293 specific, we generated a HA-NL-CD63 CRISPR knock-in pancreatic cancer (PANC-1) cell line and observed that PI4KIIIβ inhibition also reduced the bafilomycin-induced HA-NL-CD63 secretion from these cells (Supplementary Fig. 4f). Furthermore, analysis of EVs collected from control and bafilomycin-induced WT and PI4KIIIβ KO HeLa cells (Roulin et al., 2018) revealed that bafilomycin-induced EV secretion in HeLa cells also partially depends on PI4KIIIβ (Supplementary Fig. 4g).

PI4KIIIβ phosphorylates phosphatidylinositol (PI) to generate the signaling lipid phosphatidylinositol-4-phosphate (PI4P). In addition to PI4KIIIβ, the human genome encodes for three other phosphatidylinositol 4-kinases that generate PI4P. We questioned whether PI4P production by other phosphatidylinositol 4-kinases may also contribute to bafilomycin-induced EV secretion, but we did not observe any reduction in bafilomycin-induced HA-NL-CD63 secretion upon PI4KIIα or PI4KIIIα inhibition (Fig. 4i). PI4P can be further by phosphorylated phosphatidylinositol-4-phosphate 5-kinases (PIP5Ks) to generate PI4,5P2. Inhibition of PIP5K1α and PIP5K1γ, two of the three human PIP5Ks, did not affect bafilomycin-induced HA-NL-CD63 secretion (Supplementary Fig. 4h,i). These results suggest that PI4P produced specifically by PI4KIIIβ contributes to bafilomycin-induced EV secretion.

In summary, a kinase inhibitor screen revealed a role for the lipid kinase PI4KIIIβ in bafilomycin-induced exosome secretion from multiple cell lines.

### PI4KIIIβ contributes to secretory autophagy upon lysosome inhibition

Loss of PI4KIIIβ kinase activity results in a partial inhibition of bafilomycin-induced EV secretion. This raises the question as to whether PI4KIIIβ is involved in the production of a specific EV subpopulation, or contributes to all bafilomycin-induced EV secretion. To answer this question, we assessed the effect of PI4KIIIβ KO on the protein content of bafilomycin-induced EVs using mass spectrometry. Similar to the loss of SNAP23, the loss of PI4KIIIβ reduced the total amount of protein in the EV pellet and decreased the number of different proteins that could be reliably detected (Fig. 5a). However, differential abundance analysis on proteins that were present in both samples did not reveal any significant changes in the EV-associated proteomes of bafilomycin-treated WT and PI4KIIIβ KO cells (Supplementary Fig. 5). This suggests that PI4KIIIβ contributes to the production of all bafilomycin-induced EVs and does not play a major role, if any, in EV cargo selection. Moreover, the fact that PI4KIIIβ KO does not affect the ratio of autophagy cargo receptors and exosomal proteins, suggests that PI4KIIIβ plays a role in SALI. We tested this notion by quantifying the EV-associated secretion of the autophagy cargo receptor SQSTM1/P62 and could confirm that PI4KIIIβ KO inhibits SALI (Fig. 5b,c).

**Figure 5.**
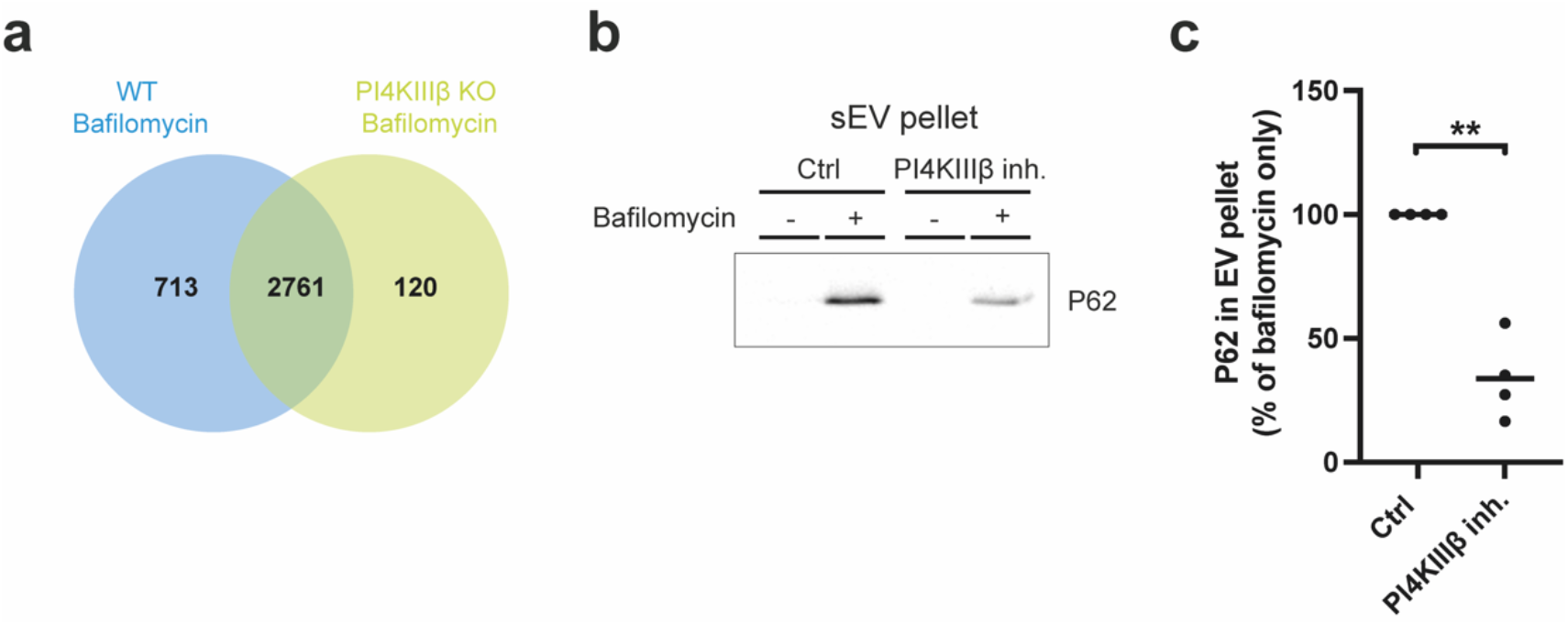
PI4KIIIβ contributes to SALI. a Venn diagram of the proteins in EVs from bafilomycin-induced wildtype (WT) and PI4KIIIβ KO HEK293 cells isolated by UF-SEC-UF. b) Western blot against P62/SQSTM1 on isolated small EVs from control and bafilomycin-induced HEK293 cells in the presence or absence of PI4KIIIβ inhibitor (PI4KIII Beta inhibitor 3, 1 μM). Representative of 4 independent experiments. c) Quantification of Fig. 5b. Horizontal bars represent the mean of four independent experiments. **P<0.01 using a one-sample, two-tailed T-test.

## Discussion

Over the last decades it has become clear that EV-mediated intercellular communication contributes to physiological homeostasis (Kalluri and LeBleu, 2020). However, dysregulation of EV-mediated communication has been demonstrated to drive disease progression in *in vivo* models of cancer (Peinado et al., 2012; Poggio et al., 2019), neurodegenerative disease (Asai et al., 2015), and osteoarthritis (Liu et al., 2021). Hence, there is a clinical need for the identification of pharmacological strategies to block distorted EV-communication in disease. In this study, we demonstrated that insertion of the small and ultra-bright luciferase NanoLuc into the CD63 gene enables robust, easy and fast quantification of EV secretion, allowing drug screens to identify modulators of EV secretion.

During the course of this study the use of NanoLuc-tagged CD63, or other EV-cargo, has been proposed in various strategies to quantify EV secretion (Cashikar and Hanson, 2019; Grisard et al., 2022; Gupta et al., 2020; Hikita et al., 2018). The approach described in this study brings together two important advances. Firstly, in contrast to several other CD63-luciferase constructs, we fused the NanoLuc enzyme to the N-terminus of CD63 in order to prevent disturbing CD63’s C-terminal internalization motif (Rous et al., 2002). Furthermore, instead of exogenous HA-NL-CD63 overexpression, we used CRISPR-genome editing to insert HA-NL into the endogenous CD63 gene. This is critical since CD63 itself drives EV secretion (van Niel et al., 2011) and overexpression may cause unwanted changes in cellular physiology. Others (Grisard et al., 2022; Gupta et al., 2020) and we (data not shown) observed that in some cell lines, such as HeLa, overexpression of NanoLuc-tagged CD63 results in cleavage and secretion of the NanoLuc-tag as free soluble protein. In this case, simply measuring NanoLuc activity in the supernatant will lead to an overestimation of EV secretion. CRISPR-mediated knock-in has some practical limitations, as it is not suitable for most primary cells and can be challenging for cells that are not easily transfected or expanded from single cells. In such cases, controlled (low-level) lentiviral exogenous HA-NL-CD63 expression as described previously (Cashikar and Hanson, 2019) may be sufficient to pick up true ‘hits’. However, we recommend that size-exclusion chromatography is performed with each generated cell line to ascertain whether the signal from EV-bound-NanoLuc secretion is not dwarfed by that of free NanoLuc.

The use of HA-NL-CD63 has several advantages compared to more traditional EV quantification techniques, such as direct particle measurements or western blot. HA-NL-CD63 allows for measurements without prior EV isolation, enables quantification of EV secretion over time frames as short as two hours, and permits EV quantification in serum-containing culture medium. Despite these advantages, our reporter only quantifies EV-associated CD63 secretion, and thus does not provide a direct quantification of EV number. Furthermore, compounds that interfere with CD63 trafficking or sorting into EVs, or compounds that directly inhibit NanoLuc enzyme activity (Grisard et al., 2022), may be picked up as false hits. Therefore, validation of compounds outside of the HA-NL- CD63 screen, using gold-standard EV isolation and quantification techniques according to MISEV criteria (Thery et al., 2018) is important.

Although CD63 is highly enriched in late endosomes, and therefore historically considered an exosome marker, recent studies have suggested that under standard in vitro culture conditions most EV-associated CD63 is secreted by vesicle budding from the PM, although this is likely to vary between cell types (Fordjour et al., 2022; Mathieu et al., 2021). In line with this, we observed that under regular 2D *in vitro* culture conditions, the majority of HA-NL-CD63 secretion by HEK293 cells occurs through mechanisms that are independent of nSmase2 and the exocytic SNARE protein SNAP23. Furthermore, basal HA-NL-CD63 secretion is sensitive to inhibition of ROCK, a kinase that has previously been implicated in vesicle budding from the PM (Li et al., 2012; Sedgwick et al., 2015). Finally, proteomic analysis revealed that, compared to bafilomycin-stimulated EVs, EVs secreted under basal conditions are enriched in proteins associated with the cell surface. Combined, these results support the hypothesis that exosome-mediated secretion accounts for only a fraction of the total EV-CD63 secretion from our HEK293 cells under basal conditions, although we cannot formally exclude CD63 secretion from a subpopulation of nSmase2- and SNAP23-independent MVBs. The high relative contribution of PM budding to HA-NL-CD63 secretion compared to MVB-derived exosomes could either result from preferential loading of HA-NL-CD63 in ectosomes, or from the low proportion of secreted exosomes to the total number of secreted EVs. We favor the latter hypothesis, as IF analysis and EM show that HA-NL-CD63, like endogenous CD63, is highly enriched in the ILVs of MVBs compared to the PM.

Even though our results suggest that exosomes account for only a fraction of the total EV secretion from our HEK293 cells under basal conditions, CD63-pHluorin imaging demonstrates that MVB-PM fusion does occur under basal conditions. Moreover, MVB-derived exosome release underlies various EV-mediated processes *in vivo*. For example, knockdown of Rab27, a small GTPase involved in docking MVBs to the PM, abrogates EV-dependent immune evasion in cancer (Poggio et al., 2019). The strong contribution of exosomes to certain EV-mediated processes *in vivo* may be due to differences in exosome and ectosome cargo, but may also indicate that the contribution of MVB-derived exosomes to the total EV secretion is larger *in vivo* than under standard 2D culture conditions *in vitro*. We therefore wished to use our HA-NL-CD63 reporter cells not only to find modulators of ectosome secretion, but also to screen for inhibitors of exosome secretion. To this end, we stimulated our reporter cells using the vATPase inhibitor bafilomycin, which blocks MVB-lysosome fusion and has been reported to enhance exosome secretion. Indeed, bafilomycin stimulation increases the secretion of HA-NL-CD63, even at low concentration (10 nM) and at early time points (2h). The mechanism by which bafilomycin stimulates exosome secretion is unclear, and there are conflicting results on the role of endosome pH neutralization in this process. While a previous study showed that neutralization of endosomal pH by NH_4_Cl or chloroquine could increase exosome secretion (Guo et al., 2017), these compounds did not affect HA-NL-CD63 secretion in a lentiviral-based reporter cell line (Cashikar and Hanson, 2019). Here we confirmed in HA-NL-CD63 endogenous reporter HEK293 cells that NH_4_Cl and chloroquine do not increase HA-NL-CD63 secretion to a similar extent as bafilomycin, suggesting the existence of a pH-independent role of vATPase in exosome secretion.

vATPase inhibition not only affects MVB-lysosome fusion, but also impairs the lysosomal degradation of autophagosomes. A recent study has demonstrated that this can result in the fusion of autophagosomes with MVBs, and subsequent exocytosis of EV-associated autophagy cargo (SALI) (Solvik et al., 2022). Our proteomics results indeed revealed a large number of autophagy proteins in the EV-pellet of bafilomycin-stimulated cells, suggesting that at least part of the bafilomycin-induced exosome release is the result of amphisome-PM fusion. Furthermore, our results confirm earlier findings that SNAP23 is involved in both MVB-fusion (Verweij et al., 2018; Wei et al., 2017) and amphisome-PM fusion (Peng et al., 2021). Importantly, we found that canonical autophagy, and therefore amphisome formation, is not strictly required for the elevated exosome secretion upon lysosomal inhibition.

By comparing the results of a kinase inhibitor screen performed under basal and bafilomycin-induced conditions, we identified PI4KIIIβ as a druggable kinase that negatively regulates basal EV secretion, but contributes to bafilomycin-induced EV secretion and SALI. PI4KIIIβ phosphorylates phosphatidylinositol to generate PI4P, a signaling phospholipid involved in the recruitment of proteins to membranes and lipid transport over membrane contact sites (MCS). Subsequent experiments using inhibitors of other PI4K isoforms and PIP5K1α/γ, suggest that it is specifically PI4P produced by PI4KIIIβ that contributes to bafilomycin-induced EV secretion. However, we cannot exclude a potential role for PI4KIIβ or PIP5K1β, for which there are currently no specific inhibitors available. PI4KIIIβ mainly localizes to Golgi membranes where it plays essential roles in Golgi formation and function (Audhya et al., 2000; Godi et al., 1999; Sciorra et al., 2005), although a fraction of PI4KIIIβ may localize to lysosomes, where it regulates tubulation and efflux of lysosomal cargo (Sridhar et al., 2013). At ER-Golgi MCS, the transport of PI4P from Golgi to ER drives the counter transport of cholesterol and ceramide to the Golgi via oxysterol-binding protein (OSBP) and ceramide transport protein (CERT), respectively. Interestingly, recent studies have found that CERT localized at ER-MVB MCS contributes to vesicle budding and EV cargo sorting under basal conditions (Barman et al., 2022; Crivelli et al., 2022). Whether CERT is involved in PI4KIIIβ-dependent exosome secretion upon lysosomal inhibition is the subject of ongoing investigation.

In this study, we engineered a bioluminescent reporter cell line that enables high-throughput screening for modulators of EV biogenesis and secretion. Our results demonstrate that this HA-NL- CD63 cell line can be used in combination with both pharmacological as well as genetic (RNAi, CRISPR KO) manipulation of EV secretion. Using the HA-NL-CD63 screening approach, we performed a kinase inhibitor screen and identified the lipid kinase PI4KIIIβ as a novel modulator of bafilomycin-induced exosome secretion and secretory autophagy. Notably, secretory autophagy and amphisome-PM-fusion have been reported in the context of cancer (Wang et al., 2019), neurological disorders (Ejlerskov et al., 2013), and viral infection (Mutsafi and Altan-Bonnet, 2018; Solvik et al., 2022; van der Grein et al., 2022). Given the druggability of PI4KIIIβ, it will be interesting to study its contribution to these pathological processes. Overall, we demonstrate the potential of HA-NL-CD63 to elucidate the molecular mechanisms underlying EV secretion and to aid the development of pharmacological strategies to target EV secretion in disease.

## Acknowledgements

We would like to thank Prof. Urs Greber (University of Zurich, Switzerland) for kindly sharing the PI4KIIIβ KO HeLa cells. We greatly acknowledge the AO|2M microscopy core platform of VU University Medical Center Amsterdam for imaging support. We greatly acknowledge the Cell and Tissue Imaging Centre (PICT-IBiSA) #Insitut Curie for electron microscopy.

This work was financed in part through the Dutch Cancer Foundation grant KWF-11308 awarded to D.M. Pegtel. M.P. Bebelman was supported by a NWO–Amsterdam Institute for Molecules, Medicines, and Systems STAR Graduate Program grant (022.005.031) and a Cancer Center Amsterdam travel grant. C. Crudden. was supported by funding from the European Union’s Horizon 2020 research and innovation programme under the Marie Skłodowska-Curie grant agreement No. 845391.

## Materials and methods Cell culture

Cells were cultured in DMEM supplemented with 10% FBS (Perbio Sciences; HyClone), 100 U/ml penicillin G, 100 mg/ml streptomycin sulfate, and 2 mM glutamine. Cell lines were cultured in a humidified atmosphere containing 5% CO_2_ at 37°C.

### Chemicals

Bafilomycin A1 (cat. no. HY-100558), GW4869 (cat. no. HY-19363), Rapamycin (cat. no. HY-10219), PI4KIII beta inhibitor 3 (cat. no. HY-15679), UNC3230 (cat. no. HY-110150), PI-273 (cat. no. HY- 103489), Chloroquine (cat. no. HY-17589A) and ISA-2011B (cat. no. HY-16937) were from Medchemexpress. U18666A (cat. no. U3633), Ionomycin (cat. no. IO634) and GSK-A1 (cat. no. SML2453) were obtained from Merck. SAR405 (cat. no. 16979) was obtained from Cayman Chemical and BAPTA-AM (cat. no. 2787) was obtained from Tocris.

### Plasmids

The generation of CD63-pHluorin in pCMV-SPORT6 was described previously in (Verweij et al., 2018). To generate the HA-nanoluc-CD63 plasmid the NanoLuc sequence was amplified using the primer pair 5’ – ATAGAATTCATGTACCCGTACGACGTCCCAGACTACGCCGTCTTCACACTCGAAGATTTC – 3’ and 5’ – ATAGGATCCCGCCAGAATGCGTTCGCACAG -3’ to add the HA-tag, and EcoRI and BamHI restriction site. The CD63 sequence was amplified using the primer pair 5’ – ATAGGATCCATGGCGGTGGAAGGAGGAATG – 3’ and 5’ – ATATCTAGACTTACATCACCTCGTAGCCACTTCTG -3’ to add the BamHI and XbaI restriction sites. The HA-Nanoluc and CD63 sequences were then cloned into EcoRI/XbaI-digested pCMV-Sport6 vector. For lentiviral transduction, HA-NL-CD63 was subcloned to the pLenti6.3/TO/V5-DEST vector (Thermo Fisher Scientific). To generate the SEC-HA-NL sequence, HA-nanoluc was amplified from HA-NL-CD63 using the primer pair 5’ – ATAGAATTCACCATGAACTCCTTCTCCACAAGCGCCTTCGGTCCAGTTGCCTTCTCCCTGGGCCTGCTCCTGGTGTTGCCTGCTGCCTTCCCTGCCCCATACCCGTACGACGTCCCAG – 3’ and 5’ - ATATCTAGAGTTACGCCAGAATGCGTTCGCAC – 3’ to add the IL6 secretion peptide, and EcoRI and XbaI restriction sites. The SEC-HA-Nanoluc was then cloned into EcoRI/XbaI-digested pCMV-Sport6 vector.

### Transfections

Plasmid transfections were performed using Lipofectamine 2000 reagent (Invitrogen), according to the manufacturer’s instruction. For CD63-pHluorin TIRF imaging, cells were transfected with 500 ng plasmid in a FluoroDish™ (World Precision Instruments) at 50–70% confluency and imaged the next day. For NanoLuc experiments, cells were transfected at 6 well scale and the next day replated to 96 well plates.

### RNA interference

Cells were transfected with ON-TARGETplus SMARTpool siRNAs (Horizon) with DharmaFECT1 transfection reagent (Thermo Scientific), according to manufacturer’s protocol. To analyze the effect of RNA interference on HA-NL-CD63 secretion, cells were replated to 96 well plates 48h after transfection. The HA-NL-CD63 assay was performed between 56h and 96h post-transfection. The following siRNAs were used: ON-TARGETplus Non-targeting Pool (D-001810-10-05), SNAP23 ON-TARGETplus SMARTpool (L-016256-00-0005), ATP6V0C ON-TARGETplus SMARTpool (L-017620-01- 0005) and PI4KB ON-TARGETplus SMARTpool (L-006777-00-0005).

### CRISPR

In order to generate the CRISPR-HA-NL-CD63 cell lines, a gRNA targeting the N-terminus of CD63 (TGCCCCAGGCCCGGCAGCCA) was cloned into the pX459 vector according to (Ran et al., 2013). For a homology-directed repair template, a linear PCR block containing the HA-NL sequence and two 100 bp homology arms flanking the insertion site at the N-terminus of CD63 was used. HEK293 and PANC- 1 cells were transfected with the gRNA plasmid and the HDR template, selected using puromycin and replated as single cells. Positive clones were identified by measuring NanoLuc activity in the supernatant and verified by PCR analysis (Fig. 1c) and Sanger sequencing.

To generate the SNAP23 KO CRISPR-HA-NL-CD63 HEK293 cells, the Alt-R™ CRISPR-Cas9 System (Integrated DNA Technologies) was used in combination with the Neon Transfection System (ThermoFisher) according to the manufacturers instruction. The gRNA sequence used was: CCUAGUCUCUGGAAA. Two days after electroporation, cells were replated as single cells. SNAP23 KO clones were identified by western blot against SNAP23 and verified by Sanger sequencing. To generate the PI4KIIIβ KO CRISPR-HA-NL-CD63 HEK293 cells, cells were transfected with the pX459 vector containing a gRNA targeting PI4KIIIβ. The gRNA sequence used was: CCACTCAACGACACTCCCGT. PI4KIIIβ KO clones were identified by western blot against PI4KIIIβ and verified by Sanger sequencing.

### Immuno-electron microscopy

HeLa cells expressing HA-NL-CD63 were fixed in 2% PFA, 0.2% glutaraldehyde in 0.1M phosphate buffer pH 7.4. Cells were then washed with phosphate buffer, embedded in 10% (wt/vol) gelatin and infused in 2.3 M sucrose. Mounted gelatin blocks were frozen in liquid nitrogen and ultrathin sections were prepared with an Ultracut FCS ultracryomicrotome (Leica). Ultrathin cryosections were labeled with anti-NanoLuc antibody (rabbit IgG, 1:250, Promega) and protein A coupled to 10nm gold particles. Samples were examined with a FEI Tecnai Spirit electron microscope (FEI Company), and digital acquisitions were made with a numeric camera (Quemesa; Soft Imaging System).

### Live TIRF microscopy

CRISPR-HA-NL-CD63 HEK293 cells were transfected with CD63-pHluorin. The following day, cells were imaged by live TIRF microscopy using a Ti-E inverted microscope setup with a 100 CFI Apochromat TIRF, oil, 1.49/0.12 mm, a/0.17 differential interference contrast objective (Nikon) and a laser bench (Roper Technologies). Cells were imaged in an imaging chamber containing 5% CO_2_ at 37°C. Images were acquired using MetaMorph software and fusion activity was quantified as described in (Bebelman et al., 2020).

### Immunofluorescence microscopy

Cells were fixed using 4% PFA (15 min, rt), blocked and permeabilized in PBS containing 0.05% saponin and 5% BSA (1 h, rt), and subsequently stained in blocking solution (1h, rt) using monoclonal antibodies against HA-tag (rat; 11867423001; Roche) and CD63 (mouse; H5C6; 556019; BD Biosciences). Alexa Fluor^®^ 594-linked anti-mouse (A11032, Thermo Fisher Scientific) and Alexa Fluor® 488-linked anti-rat (A11006, Thermo Fisher Scientific) were used as secondary antibodies. Co-staining of HA-tag and CD63 was performed sequentially to avoid cross-reactivity of anti-rat secondary with mouse anti-CD63 antibody. Cell nuclei were stained using 4′,6-diamidino-2-phenylindole (DAPI) (D9542, Sigma-Aldrich).

Confocal laser scanning microscopy was performed using a Nikon A1R+ microscope (Nikon, Tokyo, Japan) equipped with a 20x air objective or a 60 × 1.4 oil-immersion objective. Samples were irradiated using the 405, 488, and 561 nm laser lines. The 488 and 561 channels were detected with GaAsP photo-multiplier tubes (PMTs), while the 405 channel (DAPI) was detected with a regular PMT. The samples were scanned with Nikons Galvano scanner (Nikon, Tokyo, Japan) at 2048 × 2048 pixels, corresponding to a pixel size of 104 nm. NIS-Elements (Nikon, Tokyo, Japan) was used for image acquisition and denoising, and Fiji (Schindelin et al., 2012) was used for image analysis.

### HA-NL-CD63 secretion assay

CRISPR-HA-NL-CD63 HEK293 cells were seeded at a density of 6K cells per well in a white 96-well plate (CELLSTAR; 655083; Greiner bio-one). After 24h, the culture medium was replaced with fresh culture medium, typically containing kinase inhibitors or other drugs in the presence or absence of bafilomycin. After 16h, the culture supernatant was collected and cleared of dead cells and cell debris by centrifugation at 500×g for 5min. Cleared culture supernatant was transferred into a white 96 well plate (LUMITRAC™600; 655074; Greiner bio-one). To measure NanoLuc activity in both the cell and supernatant plates, Nano-Glo assay reagent, consisting of a 1:1000 dilution of Nano-Glo substrate (Promega; N1110) in buffer (100mM 2-(N-Morpholino)ethanesulfonic acid hydrate (M8250; Sigma-Aldrich), 1mM trans-1,2-Diaminocyclohexane-N,N,N′,N′-tetraacetic acid monohydrate, 150mM KCl, 1mM DL-Dithiothreitol (D0632; Sigma-Aldrich), 35mM Thiourea (T7875; Sigma- Aldrich) and 0.5% Tergitol NP-40 (NP40S; Sigma-Aldrich), pH 6.). Luminescence was measured 5 minutes after adding NanoLuc reagent using a luminescence microplate reader (Victor3, Perkin Elmer and Glomax, Promega). Cell viability assays were performed in parallel with the HA-NL-CD63 secretion assays for all treatments. To this end CRISPR-HA-NL-CD63 HEK293 cells were seeded in a transparent 96 well plate. After treatment, the culture supernatant of cells in the transparent 96 well plate was replaced by a resazurin solution (0.036 μg/ml in PBS) and incubated for 1h at 37°C. Subsequently, the fluorescence (excitation 540/35 and emission 590/20) was measured using a microplate reader (Synergy2, BioTek and Glomax, Promega).

### High-throughput kinase inhibitor screen

For the kinase inhibitor screen, a broad-spectrum kinase inhibitor library (LY-1009, MedChem Express) containing 400 kinase inhibitors was tested using the HA-NL-CD63 secretion assay. Kinase inhibitors were screened at a 1 μM concentration for 16h in the presence or absence of 100 nM Bafilomycin A1. The screen was performed in 96 well plates, each containing 80 different kinase inhibitors and 16 control wells containing 0.01% DMSO (with or without 100 nM Bafilomycin A1). Each kinase inhibitor was tested in two replicates on different plates. For every plate, HA-NL-CD63 secretion values were normalized to the average of the DMSO control wells. Cell viability was tested using the resazurin assay described above.

### Western blot

Cells lysed with RIPA buffer or isolated EVs were run on a 10% SDS gel and blotted on a nitrocellulose membrane. For detection of CD63 and CD81 SDS-PAGE was performed under non-reducing conditions. Membranes were probed with antibodies against CD63 (mouse; H5C6; 556019; BD), CD81 (mouse; JS-81; 555675; BD), SNAP23 (rabbit; 111-202; Synaptic Systems), syntenin (rabbit; ab19903, Abcam), Flotillin 1 (rabbit; D2V7J; 18634; Cell Signaling), LC3B (rabbit; D11; 3868; Cell Signaling), PI4KIIIβ (rabbit; 13247-1-AP; Proteintech), SQSTM1/P62 (rabbit; 5114T; Cell Signaling) and HSP70 (mouse; W27; sc24; Santa Cruz), and subsequently with secondary antibodies HRP-conjugated anti-Rabbit IgG (goat; 7074S; Cell Signaling) and HRP-conjugated anti-Mouse (rabbit; P0260; DAKO). Protein expression was visualized using ECL substrate (32209; Pierce) and a ChemiDoc™ MP Imaging System (Bio-Rad).

### Extracellular vesicle isolation

EVs were isolated using a combination of ultrafiltration (UF) and size exclusion chromatography (SEC). Briefly, conditioned medium was pre-cleared of debris and dead cells by differential centrifugation steps (2× 10 minutes at 500g, 2x 15 minutes at 2000g). For ultrafiltration, Centricon Plus-70 Centrifugal Filters (Merck, UFC710008) were washed with PBS according to manufacturer’s instructions. For ultrafiltration, cleared conditioned medium (60 ml) was loaded onto a filter and concentrated to 1.5 mL by centrifugation at 3000g for 15 minutes. The sample was removed from the filter by a 2-minute reverse centrifugation step at 1000g and then further purified using size exclusion chromatography. Sepharose CL/2B (GE Healthcare, 17-0140-01) in PBS was stacked in a syringe (BD, 300912) up to a 10ml column bead volume. Samples were applied and allowed to enter the column by gravity. Collection of 0.5 ml fractions was started immediately and the EV-enriched fractions 9 and 10 were pooled to 1 ml, concentrated 10x using Amicon Ultra-2 centrifugal filters (10K, Merck, UFC201024) by centrifugation for 30 minutes at 3000g, and used for western blot analysis.

### Optiprep density gradient

SEC-isolated vesicles were further purified using density-gradient centrifugation (Tulkens et al., 2020). First, a 50% w/v iodixanol solution (Optiprep, Axis-Shield, AXS-1114542) was prepared in a Tris/Sucrose/EDTA-buffer (60mM/0.25M/6mM). To prepare the density gradient, different percentages (w/v) of iodixanol in Tris/Sucrose/EDTA-buffer (10 mM/0.25M/1 mM) were layered on top of each other in thin-walled polypropylene centrifuge tubes (Beckman, 344060), starting with 40% w/v and followed by 20, 10 and 5% (w/v). EVs were loaded on top of the gradient and centrifuged for 18 hours at 100.000g, 4°C in a SW40Ti rotor (Beckman). After centrifugation, 820 μl fractions were collected and used for further analysis.

### Size Exclusion Chromatography

Culture supernatant of HA-NL-CD63 expressing cells was cleared of dead cells and cell debris by centrifugation at 500×g for 5min and 2000×g for 10min. The cleared culture supernatant was run through a column consisting of 10 ml Sepharose CL-2B beads (GE healthcare, 17-0140-01) in filtered PBS. During size exclusion chromatography with PBS as mobile phase, 26 fractions of 500 μL were collected and measured for NanoLuc activity as described above.

### LC-MS/MS Proteomic analysis - sample preparation

EVs isolated by UF-SEC-UF from control and bafilomycin-induced CRISPR-HA-NL-CD63 WT cells, bafilomycin-induced CRISPR-HA-NL-CD63 SNAP23 KO cells and bafilomycin-induced CRISPR-HA-NL- CD63 PI4KIIIβ cells (four biological replicates per group) were SpeedVac dried (Eppendorf), and processed according to the in-gel digestion protocol (Chen et al., 2011). In brief, sample was resuspended in SDS loading buffer, boiled at 98 °C for six min and ran on the 10% SDS gels (SurePAGE Bis-Tris gels from GenScript) for approximately 10 minutes at 120 V. The gels were fixed in 50% (v/v) ethanol and 3% (v/v) phosphoric acid and briefly stained with Colloidal Coomassie Blue, the sample containing lanes were sliced and cut into blocks of approximately 1 mm^3^, destained in 50 mM NH_4_HCO_3_ and 50% (v/v) acetonitrile, dehydrated using 100% acetonitrile, and rehydrated in 50 mM NH_4_HCO_3_ containing 10 μg/ml trypsin (sequence grade; Promega). After incubation overnight at 37 °C peptides were extracted and collected in a new tube, dried using a SpeedVac (Eppendorf), and stored at – 20 °C until LC-MS analysis.

### LC-MS/MS Proteomic analysis - Sample loading and run

Peptides were redissolved in 0.1% Formic acid and 100 ng was loaded onto an Evotip, and run on a 15 cm × 150 μm, 1.9 μm Endurance Column (EV1106 from EvoSep) using the Evosep One LC system with the 30 samples per day program. Peptides were electro-sprayed into the TimsTOF Pro 2 mass spectrometer and analyzed with diaPASEF (Meier et al., 2020). The MS scan was between 400-1200 m/z. The Tims settings were 1/Ko from start to end between 0.6-1.43, 26 Da precursor selection windows with 1 Da overlap, and 16 diaPASEF scans with two precursor selection windows resulting in a 1.8 s cycle time.

### LC-MS/MS Proteomic analysis - Search and data analysis

Data were analyzed using DIA-NN 1.8 with uniport UP000005640_9606 and UP000005640_9606_additional fasta. Deep learning was used to generate the in silico spectral library. Output was filtered at 0.01 FDR (Demichev et al., 2020). The Mass Spectrometry Downstream Analysis Pipeline (MS-DAP) (version beta 0.2.5.1) was used for quality control and candidate discovery (Koopmans et al., 2022). Differential abundance analysis between groups was performed on log transformed protein abundances. Empirical Bayes moderated t-statistics with multiple testing correction by False Discovery Rate (FDR), as implemented by the DEqMS functions from the limma R package, was used as was previously described (Koopmans et al., 2018). ShinyGO (Ge et al., 2020) was used for GO analysis. Venn diagrams were made using EVenn (Chen et al., 2021) based on proteins that could be detected in at least 3 out of 4 biological replicates

### Statistical analysis

Statistical analysis was performed using Prism (9.0; GraphPad Software). Data distribution was assumed to be normal, but this was not formally tested.

## Supplementary Figures

**Supplementary Figure 1.**
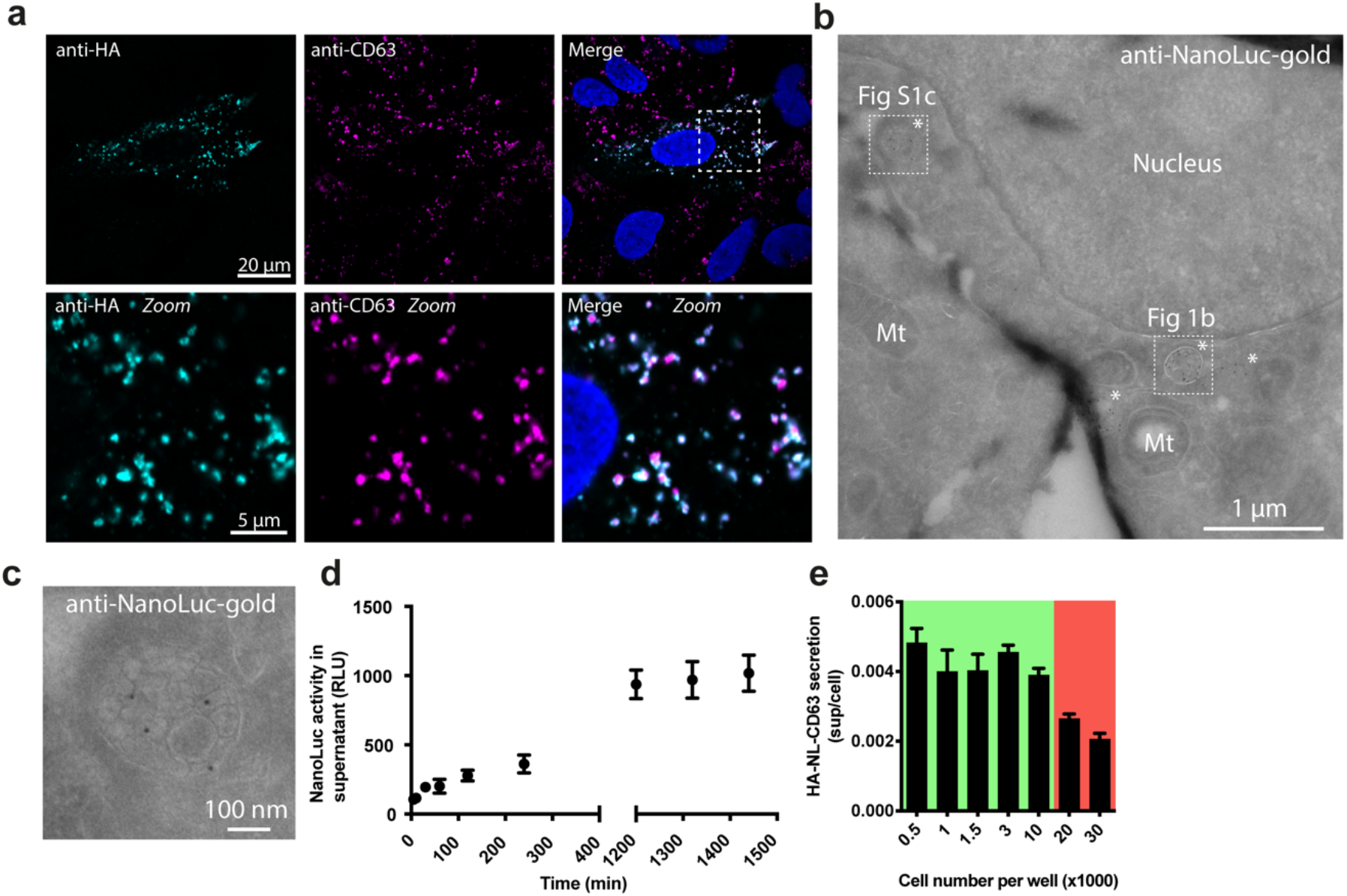
a) Immunofluorescent staining against HA (cyan) and CD63 (magenta) in HeLa cells transfected with HA-NL-CD63. Blue: DAPI. b) EM image of an HA-NL-CD63-expressing HeLa cell labelled with gold particles directed to NanoLuc (10 nm). White asterisks highlight multivesicular bodies. White squares indicate the location of the magnifications shown in Figure 1b and Supplementary Fig. 1c (S1c), right. Mt, Mitochondria. c) Zoom of Supplementary Fig. 1b. d) Increase in NanoLuc activity in the culture supernatant of HA-NL-CD63- CRISPR cells over time. Representative of two independent experiments. Data presented as mean ± S.D. e) Normalized HA-NL-CD63 secretion upon plating increasing numbers of CRISPR-HA-NL-CD63 HEK293 cells. Green and red represent sub- and over-confluent cell densities at the time of measurement, respectively. Representative of three independent experiments. Data presented as mean ± S.D.

**Supplementary figure 2.**
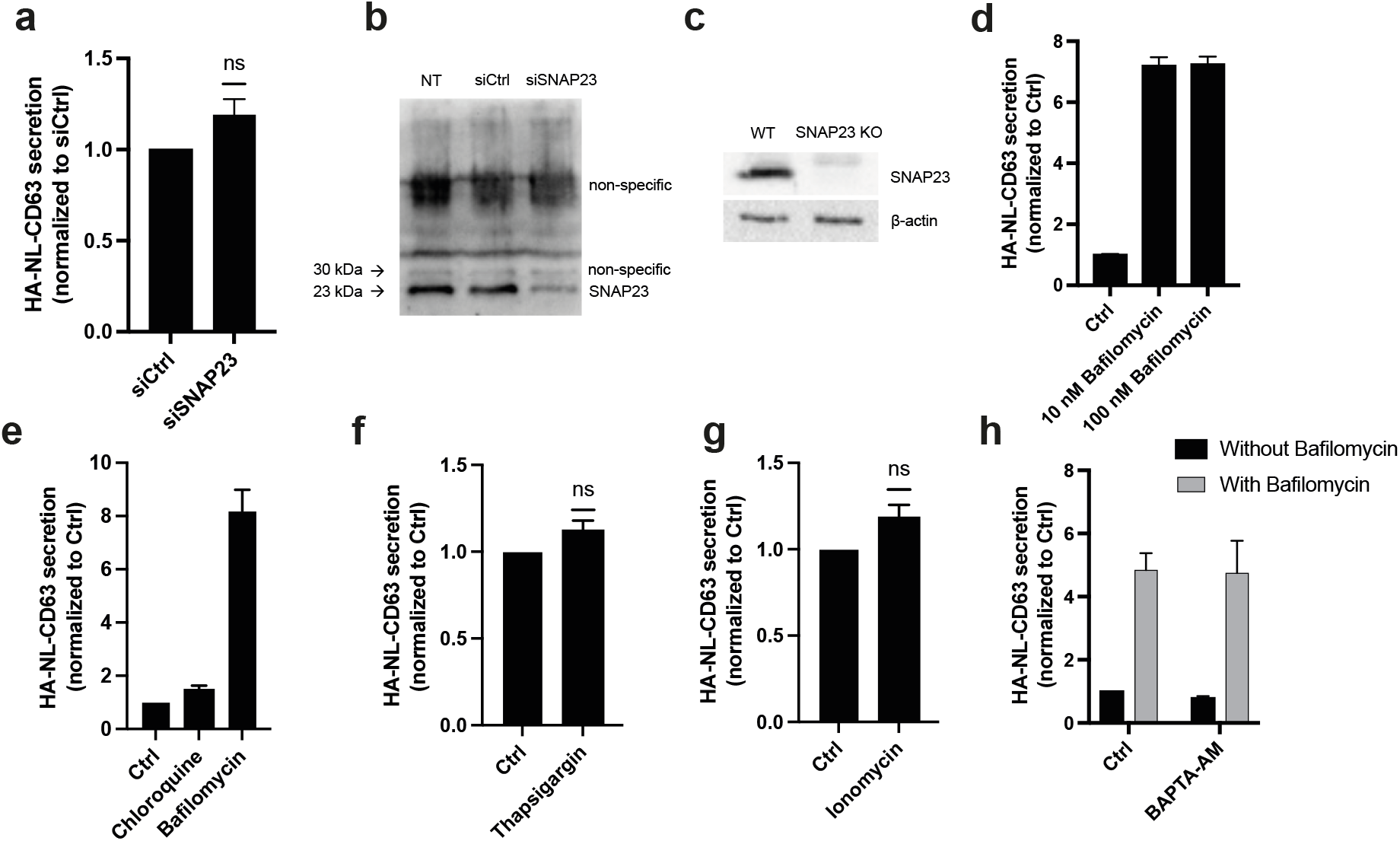
a) Effect of SNAP23 knockdown on HA-NL-CD63 secretion under basal culture conditions. Data represents the mean ± S.E.M. of seven independent experiments. ns: non-significant using a one-sample, two-tailed T-test. b,c) Confirmation of SNAP23 knockdown (b) and knockout (c) at the protein level in HA-NL- CD63-CRISPR HEK293 cells. d) Effect of 10 or 100 nM Bafilomycin on HA-NL-CD63 secretion. Representative of two independent experiments. Data represented as mean ± S.D. e-g) Effect of 50 μM chloroquine (e), 1 μM thapsigargin or 1 μM ionomycin treatment on HA-NL- CD63 secretion. Data represents the mean ± S.E.M. of three independent experiments. ns: non-significant using a one-sample, two-tailed T-test. h) Effect of 5 μM BAPTA-AM on basal and bafilomycin-induced HA-NL-CD63 secretion. Data represents the mean ± S.E.M. of three independent experiments.

**Supplementary figure 3.**
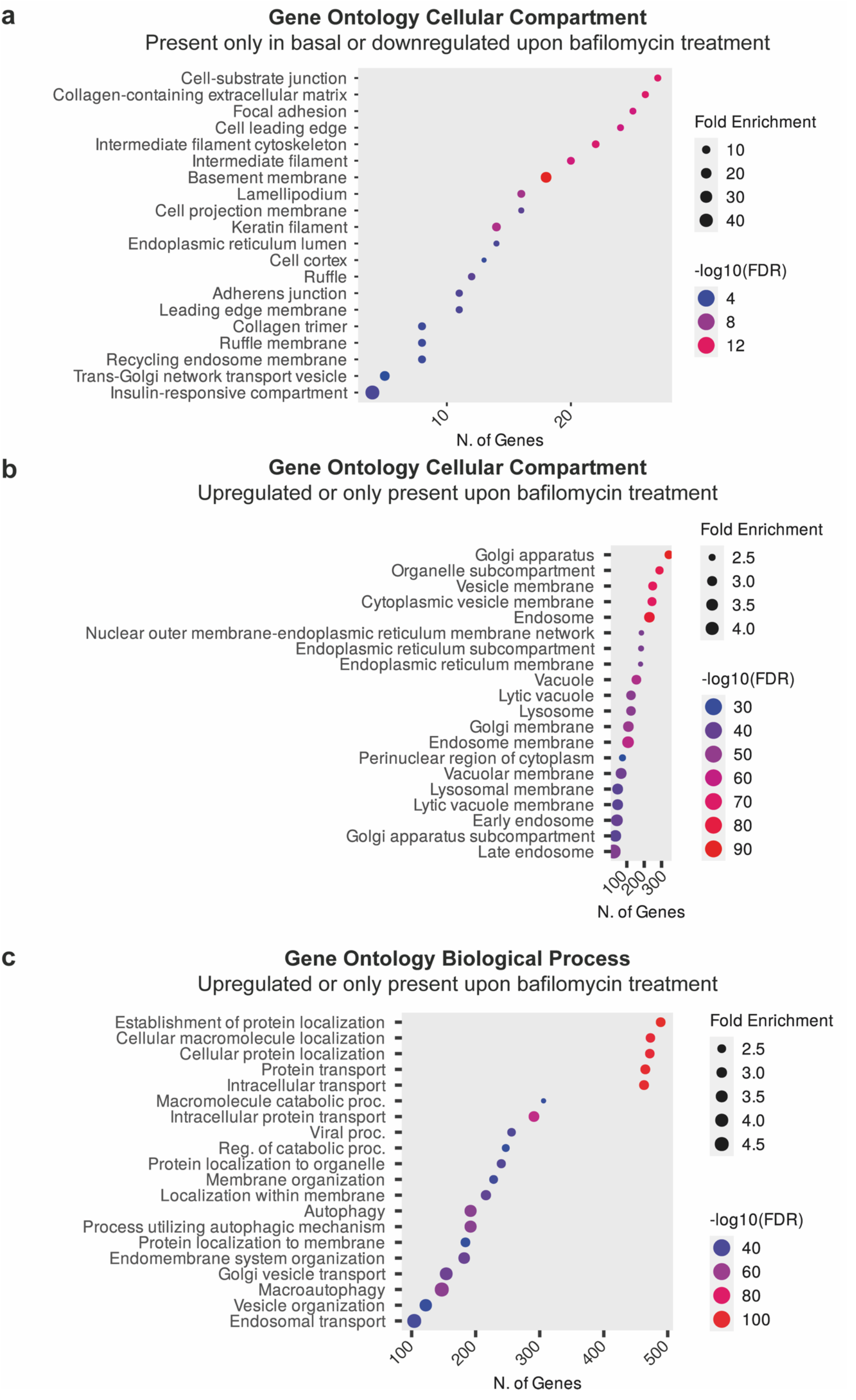
a) Gene ontology enrichment analysis of proteins that are present in EVs only under basal conditions or downregulated upon bafilomycin treatment, with the top terms for cellular component. b) Gene ontology enrichment analysis of proteins that are present in EVs only under bafilomycin-stimulated conditions or upregulated upon bafilomycin treatment, with the top terms for cellular component. c) Gene ontology enrichment analysis of proteins that are present in EVs only under bafilomycin-stimulated conditions or upregulated upon bafilomycin treatment, with the top terms for biological process. FDR: False discovery rate.

**Supplementary figure 4.**
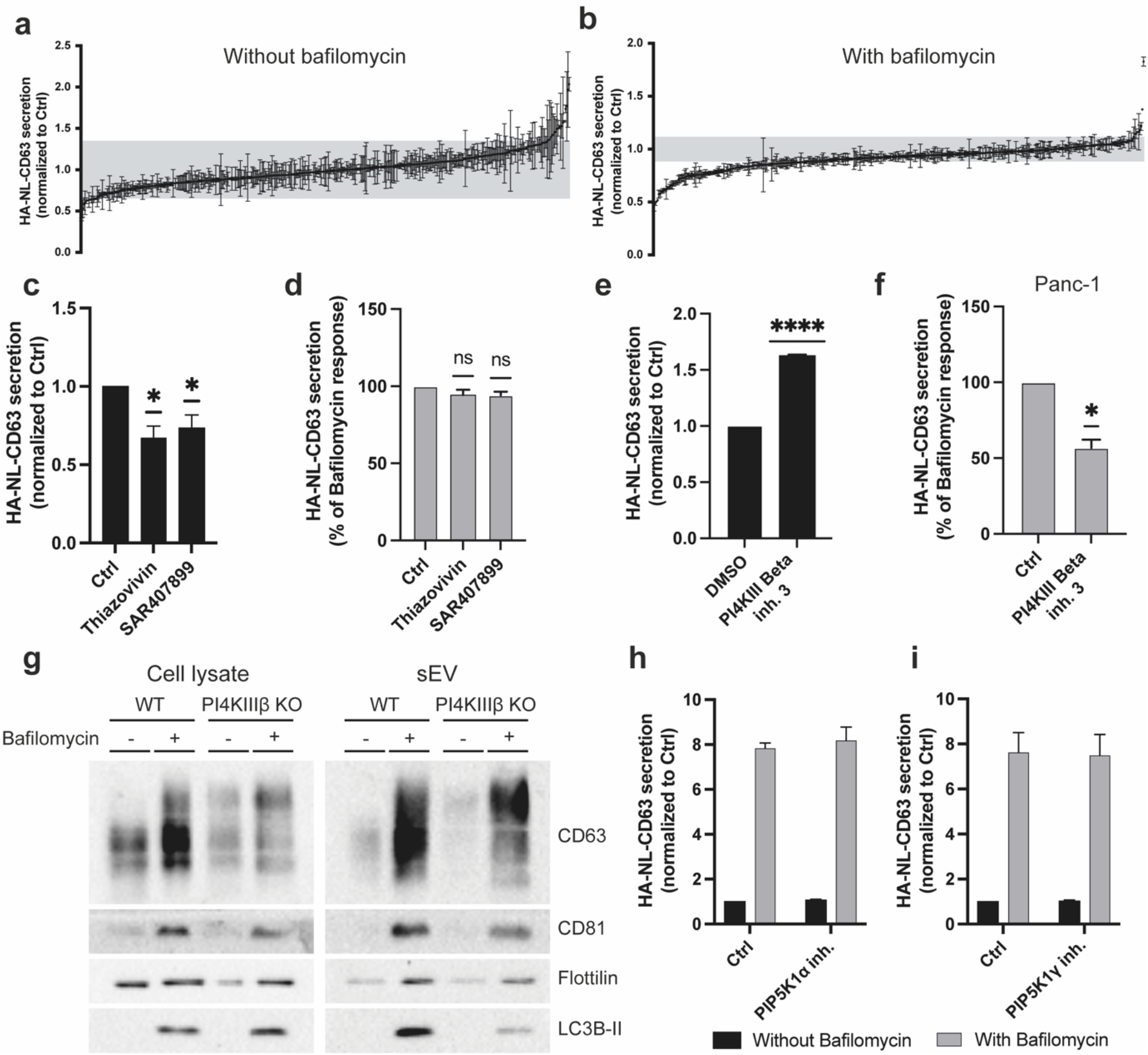
a,b) Kinase inhibitor library screens in the absence (a) or presence (b) of 100 nM bafilomycin. Data represents mean ± S.D. of two technical duplicates for each kinase inhibitor in the screens. Grey area corresponds to the mean ± 2 S.D. of the DMSO controls. c) Effect of the ROCK inhibitors thiazovivin (1 μM) and SAR407899 (1 μM) on basal HA-NL-CD63 secretion. Data represents the mean ± S.E.M. of five independent experiments. *P<0.05 using a one-sample, two-tailed T-test. d) Effect of the ROCK inhibitors thiazovivin (1 μM) and SAR407899 (1 μM) on bafilomycin-induced HA-NL-CD63 secretion. Data represents the mean ± S.E.M. of three independent experiments. ns: non-significant using a one-sample, two-tailed T-test. e) Effect of PI4KIIIβ inhibition (PI4KIII Beta inhibitor 3, 1 μM) on basal HA-NL-CD63 secretion. Data represents the mean ± S.E.M of four independent experiments. ****P<0.0001 using a one-sample, two-tailed T-test. f) Effect of PI4KIIIβ inhibition (PI4KIII Beta inhibitor 3, 1 μM) on bafilomycin-induced HA-NL-CD63 secretion from CRISPR-HA-NL-CD63 Panc-1 cells. Data represents the mean ± S.E.M of three independent experiments. *P<0.05 using a one-sample, two-tailed T test. g) Western blot against EV marker proteins on cell lysates and isolated small EVs from control and bafilomycin-induced wildtype (WT) or PI4KIIIβ KO HeLa cells. Representative of two independent experiments. h) Effect of PIP5K1α inhibition (ISA- 2011b, 20 μM) on HA-NL-CD63 secretion under basal and bafilomycin-stimulated conditions. Data represents the mean ± S.E.M. of two independent experiments. i) Effect of PIP5K1γ inhibition (UNC3230, 1 μM) on HA-NL-CD63 secretion under basal and bafilomycin-stimulated conditions. Data represents the mean ± S.E.M. of three independent experiments.

**Supplementary figure 5.**
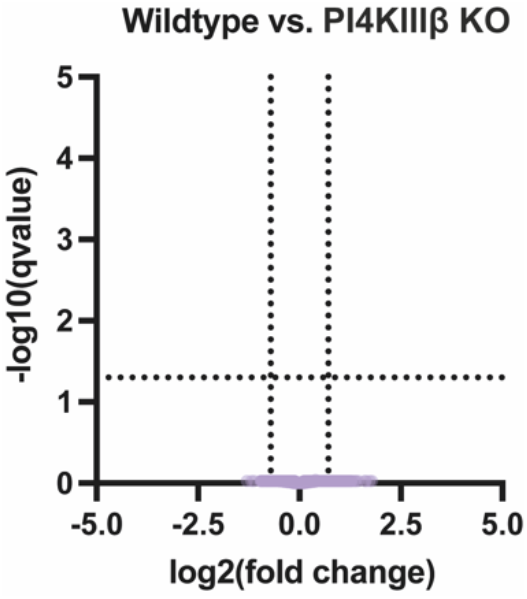
Volcano plot of the proteins in EVs from bafilomycin-induced wildtype and PI4KIIIβ KO HEK293 cells isolated by UF-SEC-UF. Proteins with a q-value ≤ 0.05, and a log2(fold change) ≤ -0.71 or ≥ 0.71 were considered differentially abundant.

## Notes

### Competing Interest Statement

The authors have declared no competing interest.

